# The cell surface hyaluronidase TMEM2 plays an essential role in mouse neural crest cell development and survival

**DOI:** 10.1101/2021.08.12.456060

**Authors:** Toshihiro Inubushi, Yuichiro Nakanishi, Makoto Abe, Yoshifumi Takahata, Riko Nishimura, Hiroshi Kurosaka, Fumitoshi Irie, Takashi Yamashiro, Yu Yamaguchi

**Author notes:** **Corresponding author:** Toshihiro Inubushi, Department of Orthodontics and Dentofacial Orthopedics, Osaka University Graduate School of Dentistry. 1-8 Yamada-oka, Suita, Osaka, 565-0871, Japan.

## Abstract

Neural crest cells (NCCs) are a migratory population that gives rise to a diverse cell lineage, including the craniofacial complex, the peripheral nervous system, and a part of the heart. Hyaluronan (HA) is a major component of the extracellular matrix, and its tissue levels are dynamically regulated in during development. Although the synthesis of HA has been shown to exert substantial influence on embryonic morphogenesis, the functional importance of the catabolic side of HA turnover is poorly understood. Here, we demonstrate that the transmembrane hyaluronidase TMEM2 plays an essential role in NCC development and the morphogenesis of their derivatives. *Wnt1-Cre*–mediated *Tmem2* knockout (*Tmem2^CKO^*) mice exhibit severe craniofacial and cardiovascular abnormalities. Analysis of *Tmem2* expression using *Tmem2* knock-in reporter mice reveals that *Tmem2* is expressed at the site of NCC delamination in the neural tube and in Sox9-positive emigrating NCCs, suggesting that *Tmem2* is critical for NCC development. Consistent with this possibility, linage tracing analysis reveals that the contribution of *Wnt1-Cre*–labeled cells to NCC derivatives is significantly reduced in a *Tmem2*-deficient background. Moreover, the emigration of NCCs from the neural tube is greatly reduced in *Tmem2^CKO^* mice. *In vitro* assays demonstrate that *Tmem2* expression is essential for the ability of mouse O9-1 NCCs to form focal adhesion on and migrate into HA-containing substrates. *Tmem2^CKO^* mice also exhibit increased apoptotic cell death in NCC-derived tissues. Collectively, our data demonstrate that *Tmem2* is essential for normal development of NCC-derivatives, including the craniofacial complex, and that TMEM2-mediated HA degradation allows NCCs to generate a tissue environment suitable for efficient focal adhesion assembly and migration. This study reveals the hitherto unrecognized functional importance of the catabolic side of HA metabolism in embryonic development and highlights the pivotal role of *Tmem2* in the process.

**Author Summary:** The functional significance of hyaluronan (HA) in embryonic developmental processes has been demonstrated by studies using genetic manipulation of HA synthesis. However, the expression of HA is regulated not only by its synthesis, but also by its degradation. This issue is of particular importance due to the extremely rapid metabolic turnover of HA. Curiously, mice with mutations/ablations of known hyaluronidase molecules, such as the lysosomal hyaluronidases HYAL1 and HYAL2, exhibit little embryonic phenotypes. This suggests the existence of yet another hyaluronidase molecule that plays a key role in regulating extracellular HA balance in developing tissues. In this context, transmembrane protein 2 (TMEM2) is a novel hyaluronidase that functions on the cell surface. Here, we demonstrate that TMEM2 is expressed at the site of neural crest development and in neural crest cell (NSC)-derived craniofacial tissue, and that NCC-targeted *Tmem2* conditional knockout mice develop severe craniofacial defects, which attests to a requirement for TMEM2-mediated extracellular HA degradation in neural crest development. Our *in vitro* and *in vivo* analyses on the underlying mechanisms of the phenotype demonstrate that TMEM2 is essential for generating a tissue environment suitable for efficient focal adhesion formation by NCCs. This paper reveals for the first time that the catabolic machinery for HA exerts a specific regulatory role in embryonic morphogenesis, and that its dysregulation of HA degradation leads to severe developmental defects.

## Introduction

Craniofacial anomalies, including midface hypoplasia and cleft lip and/or palate, account for one-third of all congenital birth defects [1]. Normal craniofacial development is an intricate biological process that requires the action of a number of distinct cell autonomous and cell non-autonomous factors and pathways. Among these, the extracellular matrix (ECM) plays a particularly important role. Cranial neural crest cells (NCCs) give rise to a major portion of the facial mesenchyme that forms the craniofacial complex, consisting of the frontonasal process and the first pharyngeal arch. NCCs are a migratory population of cells that arise at the edge of the neural tube during neurulation, and contribute to the formation of a variety of tissues, including the craniofacial tissues, cardiovascular system, peripheral nervous system and skeleton [2]. After induction and specification at the edge of the neural tube, NCCs undergo a process called delamination, in which they emigrate from their site of origin and subsequently migrate toward target sites. Mutations in genes involved in NCCs development can lead to a wide range of human congenital malformations, including craniofacial anomalies.

Hyaluronan (HA) is a non-sulfated glycosaminoglycan widely distributed in the ECM of a variety of tissues. The importance of HA in the development of NCC-derived tissues is illustrated by the observation that disruption of HA synthesis by means of genetic ablation of the *Has* genes, which encode hyaluronan synthases, leads to defects in NCC-derived tissues. In mice, *Has2* ablation results in mid-gestation (E9.5-10) lethality due to severe defects in endocardial cushion formation [3, 4]. Defects in these cardiac structures are often associated with the abnormal development of a subpopulation of cranial NCCs [5]. Cranial NCC-targeted conditional *Has2* knockout mice also exhibit these defects in formation of the NCC-derived craniofacial structures [6]. In zebrafish, mutations in hyaluronan synthase genes (*has1* and *has2*) result in impairments in NCCs migration and craniofacial defects [7].

Tissue levels of ECM molecules are controlled by the balance between synthesis and degradation. A unique feature of HA among ECM molecules is its extremely rapid turnover [8–10]. An estimated one-third of the total body HA (∼15 g in a person with a 70 kg body weight) is degraded and turned over daily [9], and the half-life of HA in skin is only 1 to 1.5 days [8]. Although the rate of HA degradation in embryonic tissues has not been determined, it seems reasonable to speculate that HA is actively degraded in embryonic tissues, since HA staining changes dynamically in intensity during embryonic development [11–13]. Although several classes of hyaluronidase proteins have been identified, the identity of cell surface (or secretory) hyaluronidase(s) that is directly involved in the degradation of extracellular HA has remained elusive. The HYAL family proteins, including HYAL1, HYAL2, and SPAM1, have been extensively studied as principal hyaluronidases in mammalian species. However, while some data suggest that a subpopulation of HYAL proteins is present on the cell surface [14, 15], their physiological sites of action have been controversial. In fact, multiple lines of evidence indicate that HYAL proteins are primarily associated with lysosomes and related intracellular vesicles [16–21]. Also, HYAL1 and HYAL2 favor acidic pH (pH < 5) for their hyaluronidase activity [19, 22], consistent with the notion that they are primarily involved in the ultimate lysosomal degradation of internalized HA fragments in lysosomes. Moreover, mice with mutations in HYAL proteins exhibit only mild developmental phenotypes. *Hyal1* null mice are viable and exhibit few developmental defects [23]. *Hyal2* null mice on a C57Bl/6 background are also viable and exhibit only mild developmental abnormalities, including shortening of the nose and widened interorbital space, whereas on a mixed C129;CD1;C57Bl/6 background, approximately half of adult *Hyal2* null mice exhibit moderate heart defects, including expanded heart valves and cardiac hypertrophy [24, 25]. These relatively mild developmental phenotypes resulting from knockout of HYAL family hyaluronidases suggests that there may be another hyaluronidase responsible for the prominent role of HA degradation during embryonic development.

Transmembrane protein 2 (TMEM2; gene symbol CEMIP2) was originally identified as a large type II transmembrane protein with an unknown function. Even in the absence of a defined function, zebrafish *tmem2* mutants (*frozen ventricles* and *wickham*) were nevertheless found to exhibit a developmental heart phenotype related to endocardial cushion defect, which is accompanied by excessive accumulation of HA [26, 27]. Subsequently, we demonstrated that TMEM2 is a hyaluronidase that degrades extracellular high-molecular weight HA into intermediate-sized HA fragments at near neutral pH [28]. In mouse embryos, *Tmem2* is expressed in a developmentally regulated manner, with the peak of expression prior to E11 and with prominent sites of expression in the neural tube, the first branchial arch, and the frontonasal processes [29]. More recently, we have demonstrated that TMEM2 plays a critical role in promoting integrin-mediated cell adhesion and migration via its removal of anti-adhesive HA from focal adhesion sites [30]. These observations suggest to us that TMEM2 is the key hyaluronidase that regulates dynamic remodeling of the HA-rich ECM during NCCs development and migration.

In this report, we use *Wnt1-Cre*–mediated conditional ablation of *Tmem2* to determine the role of this cell surface hyaluronidase in cranial NCCs development and embryonic morphogenesis. We show that ablation of *Tmem2* leads to severe craniofacial defects, attesting to the essential role of TMEM2-mediated HA degradation in NCCs development and survival. The essential role of TMEM2-dependent HA degradation in NCCs development and craniofacial morphogenesis provides initial insight into the critical importance of dynamic HA turnover and matrix remodeling in embryonic development.

## Results

### Wnt1-Cre mediated conditional *Tmem2* knockout results in multiple defects in craniofacial development

To gain an initial insight into the role of TMEM2 during NCCs development, we used *in situ* hybridization to examine the spatiotemporal expression pattern of *Tmem2* during the mid-gestation period (E8.5-10.5). In whole mount preparations at E8.5 and 9.0, expression of *Tmem2* is detected in the neural tube, frontonasal region, branchial arches, and heart (Fig 1A, *Dorsal, Lateral*). Transverse sections of the E9.0 neural tube further demonstrate that *Tmem2* expression in the neural tube is concentrated in the dorsal part (Fig 1A, *Transverse*), from which NCCs arise. At E9.5 and E10.5, expression of *Tmem2* is detected in the forebrain, midbrain, hindbrain, trigeminal ganglion, branchial arches, heart, and dorsal root ganglia (**S1 Fig**). Thus, *Tmem2* expression is associated with tissues involved in neural crest development.

**Fig 1.**
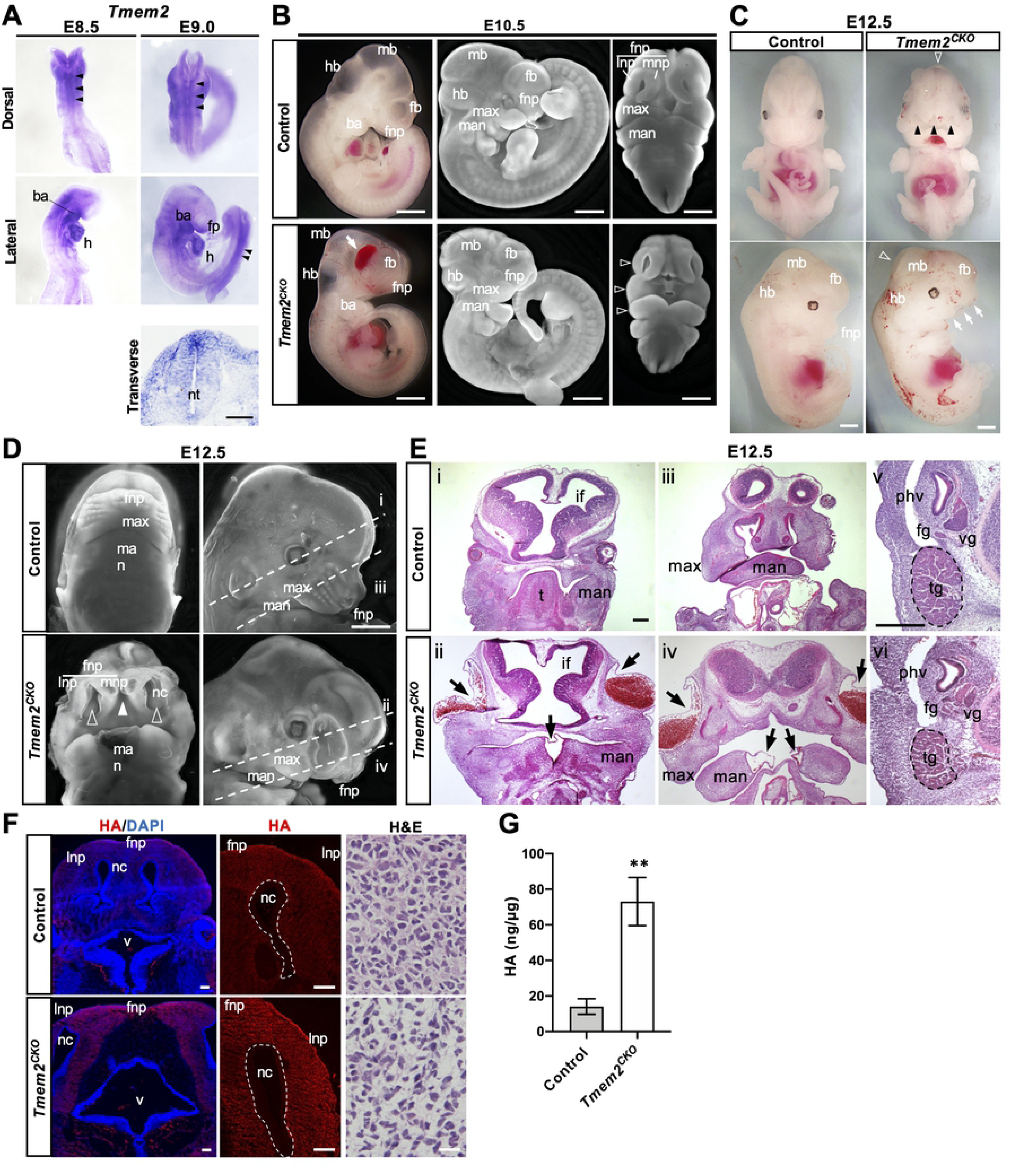
NCC-targeted conditional knockout of *Tmem2* leads to craniofacial malformations. **(A) Expression of *Tmem2* in wild-type embryos by *in situ* hybridization.** Whole-mount images of the lateral and dorsal aspects of embryos and a transverse section through the neural tube at E8.5 and E9.0. Robust *Tmem2* expression is observed in the dorsal midline region of the neural tube (*arrowheads*), as well as in the facial prominence (*fp*), the branchial arches (*ba*), heart (*h*), neural tube (*nt*). (**B**) **Gross phenotype of *Tmem2^CKO^* embryos at E10.5.** Whole-mount images of unstained embryos (*left panels*) and fluorescence microscopic images of DAPI-stained embryos (*center* and *right panels*) are shown. Hypoplasia of the frontonasal process , and maxillary process and mandibular process, is observed in *Tmem2^CKO^* embryos (*open arrowheads*). *Arrow* indicates edema in the facial region. (**C**) **Gross phenotype of *Tmem2^CKO^* embryos at E12.5.** Whole-mount images reveal severe developmental malformations of the craniofacial region of *Tmem2^CKO^* embryos with hypoplasia of the facial processes (*arrows*) and the lack of fusion of the frontonasal process (*filled arrowheads*). Mild exencephaly-like defects are also found in *Tmem2^CKO^* embryos (*open arrowhead*s). Hemorrhagic lesions are also frequently observed in *Tmem2*^CKO^ embryos. **(D, E) Phenotypes in the craniofacial region of E12.5 *Tmem2^CKO^* embryos.** (**D**) Fluorescence microscopic images of the craniofacial region of DAPI-stained embryos at E12.5. *Tmem2^CKO^* embryos show lack of fusion of the bilateral medial nasal processes (*filled arrowhead*) and bilateral mandible processes in the midline, lack of fusion between frontonasal process and maxillary process (*open arrowheads*), wide opening nasal cavity, and noticeably wider heads. Broken lines (i, ii, iii and iv) indicate the orientations of sections shown in *E*. (**E**) H&E staining of transverse sections through the forebrain (i, ii) and the maxillary region (iii, iv). (v, vi) High magnification images corresponding to the area of NCC-derived peripheral nervous tissues, including trigeminal (*tg*), vestibular (*vg*), and facial (*fg*) ganglia. Note that *Tmem2^CKO^* embryos exhibit a reduction in the size of these NCC-derived ganglia. *Tmem2^CKO^* embryos also exhibit blister-like epithelial detachment, which is often accompanied with hemorrhaging, in the lateral portion of the frontonasal process and at the midline of the mandibular arches (*arrows*). **(F) Accumulation of HA in craniofacial tissues of E12.5 *Tmem2^CKO^* embryos.** Transverse sections of the facial processes were double-labeled with bHABP and/or DAPI (*left* and *center panels*) or stained with H&E (*right panels*). Increased HA staining is observed in the facial processes of *Tmem2^CKO^* embryos (*HA*). H&E staining reveals expanded intercellular spaces in *Tmem2^CKO^* embryos (*H&E*). **(G) Quantification of HA in the tissue lysates from the facial tissues of E12.5 embryos.** Data represent means ± SD (*n* = 5). \*\**P* < 0.01 by Student’s *t*-test. ns, not significant. *ba*, branchial arch; *h*, heart; *fnp*, frontonasal process; *fp*, facial prominence; *max*, maxillary process; *man*, mandibular process; *fb*, forebrain; *mb*, midbrain; *hb*, hindbrain; *nc*, nasal cavity; *nt*, neural tube; *mnp*, medial nasal process; *lnp*, lateral nasal process; *t*. tongue; *phv*, primary head vein; *tg*, trigeminal ganglion; *vg*, vestibular ganglion; *fg*, facial ganglion. Scale bars: 250 μm in **A**, **F** (*left panels*); 500 μm in **B**-**E**; 25 μm in **F** (*right panels*).

The *Wnt1-Cre* driver has been used to determine the function of genes in NCCs development, migration, and subsequent differentiation [31, 32]. To determine the role of *Tmem2* in the development of NCCs and their derivatives, we generated a conditional *Tmem2* allele (*Tmem2^flox^*; see **Materials and Methods**) and crossed *Tmem2^flox/flox^* mice with *Wnt1-Cre* mice. While heterozygous *Wnt1-Cre;Tmem2^flox/wt^* mice were born alive without detectable developmental defects and were fertile, no homozygous conditional mutants (i.e., *Wnt1-Cre;Tmem2^flox/flox^*; hereafter referred to as *Tmem2^CKO^*, hereafter) were recovered at birth. Therefore, we performed timed matings to obtain homozygous embryos at several time points between E10.5 and E12.5. At E10.5, *Tmem2^CKO^* embryos exhibit hypoplasia of the frontonasal, maxillary, and mandibular processes (Fig 1B). Hemorrhage and edema were frequently observed in the craniofacial regions of *Tmem2^CKO^* embryos at times later than E10.5 (Fig 1B). In addition, the growth retardation is often observed in these mutant embryos at this stage. At gestational stages later than E12.5, *Tmem2^CKO^* embryos exhibit severe craniofacial abnormalities (42 of 42 embryos, 100%), characterized by reduced outgrowth of the frontonasal and maxillary processes, lack of fusion of the bilateral medial nasal processes and bilateral mandible processes at the midline, lack of fusion between frontonasal process and maxillary process, wide opening nasal cavity, and noticeably wider heads (Fig 1C, 1D). Histomorphological analysis of E12.5 *Tmem2^CKO^* embryos further demonstrates the hypoplastic but laterally expanded maxillary components and the lack of fusion of the facial processes (Fig 1E-ii, iv). Epithelial blistering was frequently observed in the lateral portion of the frontonasal process and in the midline region of the mandibular arches (34 of 42 embryos, 81.0%) (*arrows* in Fig 1E-ii, iv). Defects in branchial arch derivatives, such as the tongue, are also consistently observed in *Tmem2^CKO^* embryos (Fig 1E-ii). NCC-derived peripheral nervous tissues, such as trigeminal, facial, and vestibular ganglia are smaller in *Tmem2^CKO^* embryos compared with control embryos (Fig 1E-vi). In addition, mild neural tube closure defects are detected in a fraction of *Tmem2^CKO^* embryos (4 of 42 embryos, 9.5%) (Fig 1C**, S2 Fig**). At E13.5, *Tmem2^CKO^* embryos could be identified, but all of them were dead and exhibited severe craniofacial and cardiovascular abnormalities.

A subpopulation of cranial NCCs, referred to as cardiac NCCs, migrates into the third, fourth and sixth branchial arches and gives rise to the aortic and pulmonary trunk, the cap of the intraventricular septum, the developing outflow tract cushions, and the parasympathetic system of the heart [33]. *Wnt1-Cre* is also active in cardiac NCCs [34]. Consistent with the functional role for TMEM2 in cardiac NCCs, *Tmem2^CKO^* embryos exhibit expanded endocardial cushions with sparse mesenchyme and absence of the aorticopulmonary septum in the outflow tract region (**S3 Fig**). The lack of endocardial cells in conotruncal endocardial cushions is indicative of defective endocardial cell migration in *Tmem2^CKO^* embryos (**S3 Fig**).

To determine whether TMEM2 indeed functions as a hyaluronidase in tissues that exhibit morphogenetic defects, we examined whether HA is increased in these tissues. HA staining using biotinylated HA binding protein (bHABP) reveals increased HA the frontonasal process of E12.5 of *Tmem2^CKO^* embryos (Fig 1F). Similar increase in HA staining is observed in the mandible and the outflow tract endocardial cushions of the heart (**S4 Fig**). Increase in tissue HA contents is also shown by quantification of HA in lysates of craniofacial tissues (Fig 1G). Hematoxylin and eosin (H&E) staining of the frontonasal process reveals that the size of extracellular spaces is increased in *Tmem2^CKO^* embryos (Fig 1F), a tissue phenotype generally associated with excess HA accumulation [35]. Altogether, these observations on *Tmem2^CKO^* embryos demonstrate that *Tmem2* is required for normal cranial neural crest development, acting as a functional hyaluronidase.

### Expression of TMEM2 in developing tissues in relation to the distribution of HA

To gain insight into the cellular mechanisms underlying the phenotype of *Tmem2^CKO^* embryos, we compared the detailed spatiotemporal expression of TMEM2 protein in relation to the distribution of HA in NCCs and their derivatives. To facilitate this analysis, we created by using the CRISPR-Cas9 gene-editing technology a knock-in mouse line (referred to as *Tmem2-FLAG^KI^* mice), in which a FLAG-tagged TMEM2 is expressed under the control of the endogenous *Tmem2* promoter (see **Materials and Methods** and **S5 Fig**). Mice homozygous for this knock-in allele are born and grow normally, indicating that TMEM2 produced from this FLAG-tagged allele is functional.

Analysis of *Tmem2-FLAG^KI^* embryos at E11.0 reveals that TMEM2 protein is expressed in the neuroepithelia of the forebrain, midbrain, and hindbrain, as well as facial prominence, branchial arches, dorsal root ganglia, and the developing heart (Fig 2A,B). These results are in good agreement with the expression of *Tmem2* mRNA detected by *in situ* hybridization (see **S1 Fig**) and confirm that TMEM2 protein is expressed in tissues populated by NCCs. Double-staining of TMEM2 and HA using anti-FLAG antibody and bHABP, respectively, reveals that TMEM2 protein and HA show a roughly complementary distribution (Fig 2A,B), supporting a role of TMEM2 protein as a functional hyaluronidase. Significantly, the sites of TMEM2 expression coincide with the sites where defects are observed in *Tmem2^CKO^* embryos, suggesting the requirement of functional TMEM2 protein in the normal development of NCC-derived tissues.

**Fig 2.**
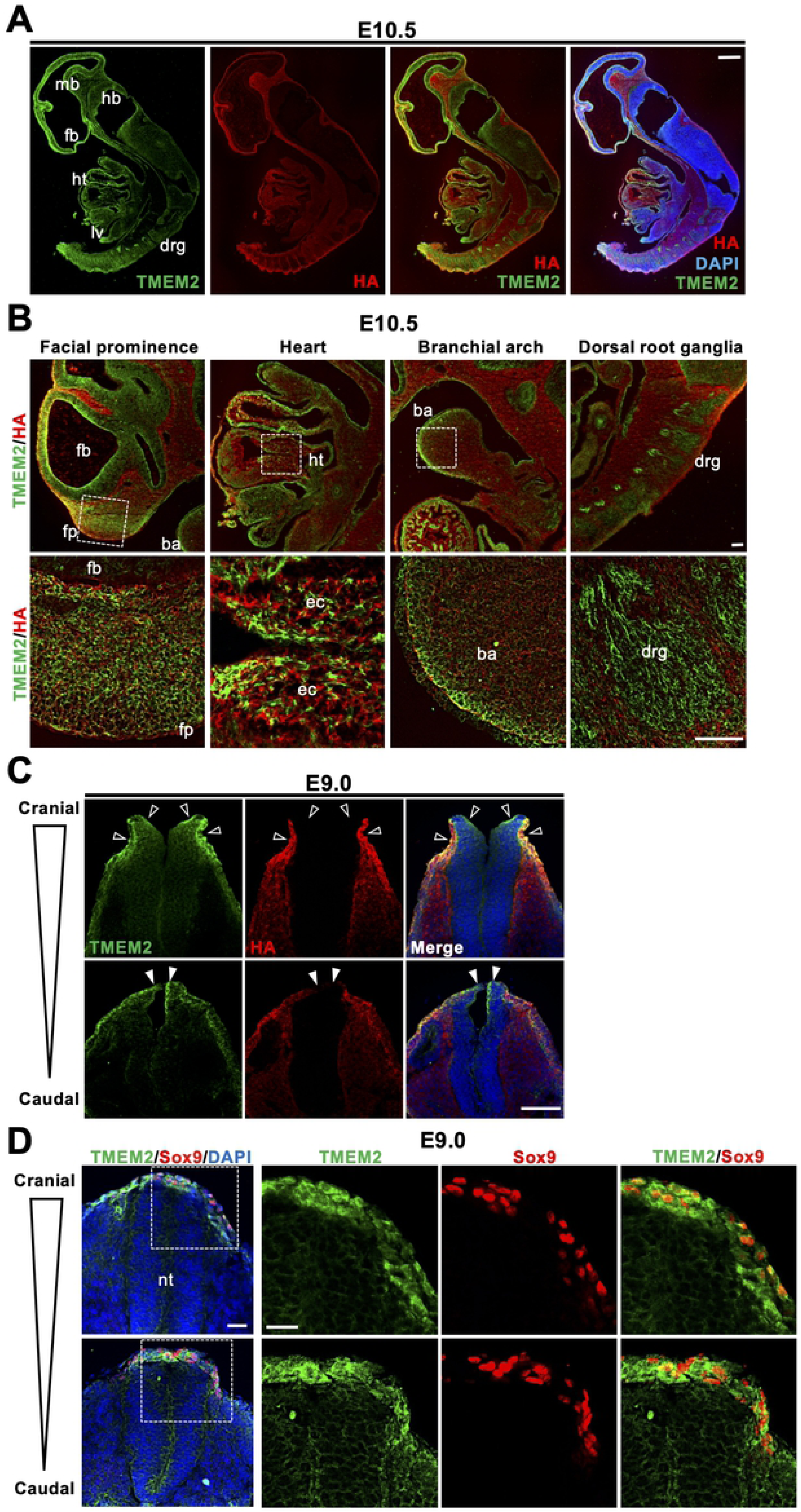
Expression of TMEM2 in relation to HA distribution in the mouse embryo. (**A**) Sagittal sections of *Tmem2-FLAG^KI^* reporter embryos were double-labeled with anti-FLAG antibody (to detect TMEM2-FLAG protein) and bHABP (to detect HA). Nuclei were counterstained with DAPI. Strong TMEM2 protein expression is observed in the neuroepithelium, facial prominence, heart, and dorsal root ganglia. *fb*, forebrain; *mb*, midbrain; *hb*, hindbrain; *ht*, heart; *lv*, left ventricle; *drg*, dorsal root ganglia. (**B**) High magnification images of the facial prominence, heart, branchial arch, and dorsal root ganglion in *Tmem2-FLAG^KI^* embryos double-labeled for TMEM2-FLAG protein and HA. Areas indicated by boxes are enlarged in lower panels. *fb*, forebrain; *fp*, facial prominence; *ba*, branchial arch; *ht*, heart; *ec*, endocardial cushion; *drg*, dorsal root ganglia. (**C**) Transverse sections of the neural tube of *Tmem2-FLAG^KI^* embryos at E9.0. Sections at the cranial and trunk levels of the neural tube were double-labeled for TMEM2-FLAG protein and HA as in *A*. High expression of TMEM2 is observed in the neural plate and the border region of the neural tube (*open arrowheads*). HA is not detectable in the dorsal midline region (*filled arrowheads*), where the pre-migratory neural crest is located. (**D**) Double-labeling of neural crest cells for TMEM2-FLAG and Sox9. Transverse sections of E9.0 neural tube were stained for TMEM2-FLAG and Sox9. Note that Sox9-positive pre-migratory and emigrating NCCs at the edge of the neural tube highly express TMEM2 on their surface. *nt*, neural tube. Scale bars: 250 μm in **A**; 50 μm in **B**; 300 μm in **C**; 100 μm in **D.**

### Role of TMEM2 in NCCs emigration

Based on the *in situ* hybridization data showing prominent expression of *Tmem2* in the neural tube (Fig 1), we examined in *Tmem2-FLAG^KI^* mice the expression pattern of TMEM2 protein during neural tube closure and neural crest formation. Distribution of HA in the pertinent regions was also examined by double-labeling with bHABP. In E9.0 embryos, TMEM2 protein is expressed in the neural fold and the neural plate border, while HA staining is absent in these regions (Fig 2C), indicating that TMEM2 functions as a hyaluronidase in these regions. Next, we analyzed the localization of NCCs by double-labeling for TMEM2 protein and Sox9, an NCCs specifier commonly used for a marker for pre-migratory and emigrating NCCs [36]. As shown in Fig 2D, Sox9-positive cells in the dorsal neural tube and those emigrated out of the neural tube highly express TMEM2 protein. These observations suggest that TMEM2 plays a role in NCCs emigration from the neural tube.

To further characterize the functional role of TMEM2 in NCCs emigration, we examined the localization of NCCs in *Tmem2^CKO^* and control embryos using Sox9 and Sox10 as markers for pre-migratory and migrating NCCs, respectively (Fig 3). In wild-type embryos at E9.0, Sox9-positive cells exhibit a normal pattern of emigration out of the neural tube (Fig 3A, *Sox9/Control*). The extracellular space surrounding these pre-migratory NCCs are devoid of HA (*open arrowheads* in *HA/Control*), indicating TMEM2-dependent removal of HA in the pericellular space of pre-migratory NCCs. In *Tmem2^CKO^* embryos, in contrast, few Sox9-positive cells are localized outside of the neural tube (Fig 3A, *Sox9/Tmem2^CKO^*) and HA staining persists in the dorsal midline region of the neural tube (*filled arrowheads* in *HA/Tmem2^CKO^*). Meanwhile, Sox10 staining (Fig 3B) demonstrates that the number of NCCs migrating along the ventral migration pathway is reduced in *Tmem2^CKO^* embryos (Fig 3B, *Tmem2^CKO^*). Counting of pre-migratory NCCs within the neural tube (Fig 3C, *upper graph*) and migrating NCCs along the ventral migration pathway (Fig 3C, *lower graph*) confirms these observations. Together, these results suggest that functional TMEM2 is required for the efficient emigration of NCCs out of the neural tube. Given the even distribution of Sox10-positive NCCs along the migration pathway (Fig 3B), it is likely that migration *per se* is not significantly affected by loss of TMEM2, and that the reduced number of migrating NCCs in *Tmem2^CKO^* embryos is mainly due to the reduced emigration of NCCs from the neural tube.

**Fig 3.**
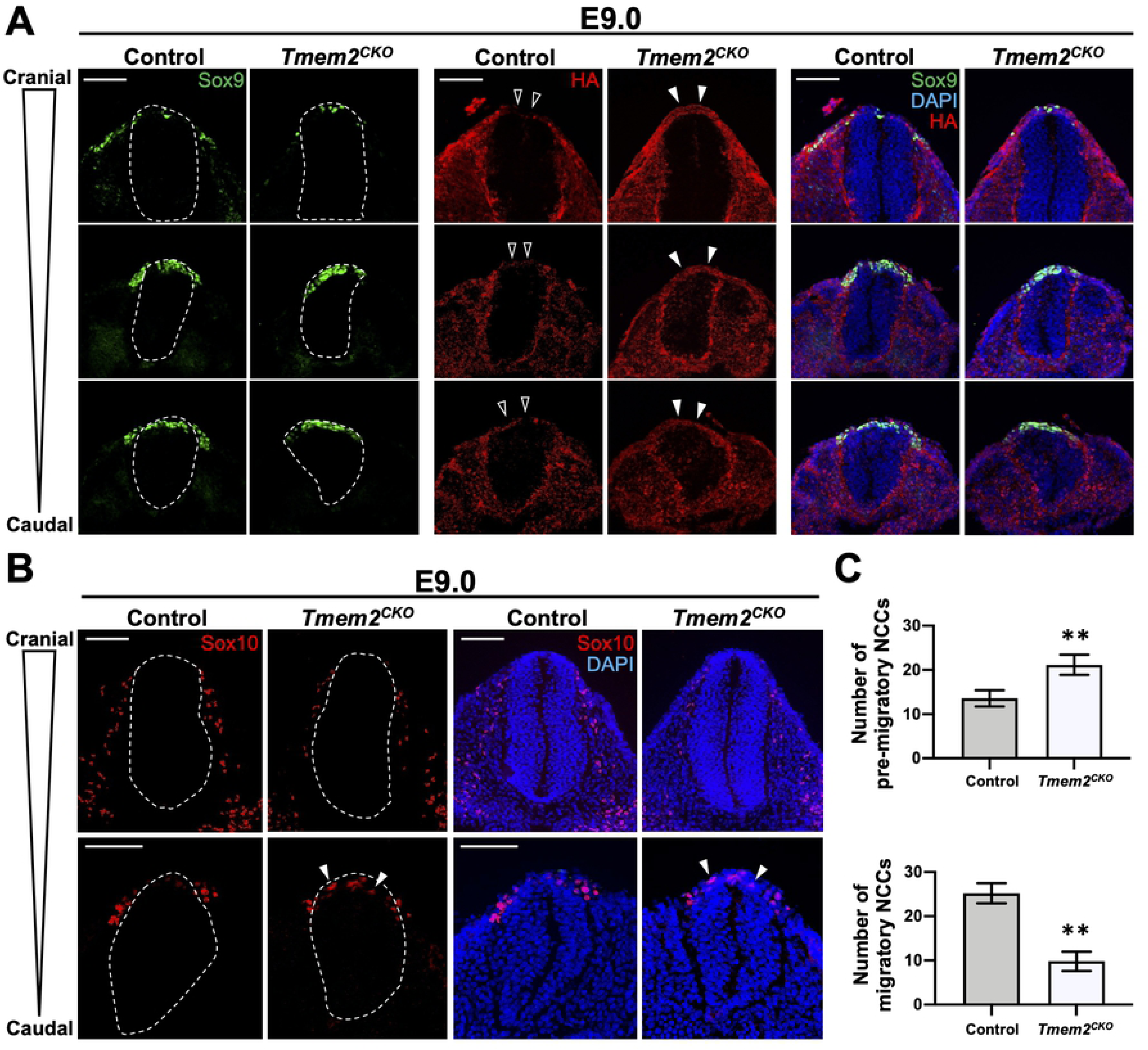
Effects of *Tmem2* ablation on NCCs development. (**A**) Transverse sections at three different levels of the neural tube of *Tmem2^CK^*^O^ and control embryos at E9.0 were triple-stained for Sox9 (*green*), HA (*red*), and nuclei (*blue*). *Broken lines* indicate the external contour of the neural tube. Note that emigration of Sox9-positive cells out of the neural tube is inhibited in *Tmem2^CK^*^O^ embryos. Also, in *Tmem2^CK^*^O^ embryos, the dorsal surface of the neural tube exhibits HA staining (*filled arrowheads*), whereas little HA staining is detectable in the corresponding area of control embryos (*open arrowheads*). (**B**) Transverse sections of *Tmem2^CK^*^O^ and control embryos at E9.0 were double-stained for Sox10 (*red*) and nuclei (*blue*). In *Tmem2^CKO^* embryos, the number of Sox10-positive migrating NCCs is significantly decreased. The image in the caudal neural tube reveals that some Sox10-positive cells persist within the neural tube in *Tmem2^CKO^* embryos (*arrowheads*). (**C**) Quantification of pre-migratory and migrating NCCs in *Tmem2^CKO^* and control embryos. The number of Sox9-positive cells and Sox10-positive cells in respective ROIs (see *Materials and Methods*) was counted as pre-migratory and migrating NCCs, respectively. Data represent means ± SD (*n* = 5) shown as horizontal bars. \*\**P* < 0.01 by Student’s *t*-test. ns, not significant. Scale bars: 300 μm in **A** and **B**.

### Lineage trancing of NCCs in *Tmem2^CKO^* mice

We next performed fate mapping experiments to determine the contribution of *Tmem2*-deficient NCCs to the formation of NCC-derived structures. For this, we crossed the ZsGreen (Z/EG) reporter gene [37] onto the *Tmem2^CKO^* background to generate *Tmem2^CKO^;Z/EG* mice. Distribution patterns of Z/EG-positive NCCs were analyzed between these and control (*Wnt1-Cre;Tmem2^wt/wt^;Z/EG*) mice. While the gross distribution pattern of green Z/EG signals in neural crest derivatives is not significantly altered, both the spatial extent and the intensity of Z/EG signals in the facial prominences and the branchial arches are reduced in the *Tmem2^CKO^* background compared to the control background at E9.5 (*open arrowheads* in Fig 4A). Meanwhile, aberrant accumulation of Z/EG-positive cells was observed in the midbrain and hindbrain of *Tmem2^CKO^;Z/EG* embryos (*filled arrowheads* in Fig 4A). Sections through the frontonasal process and the midbrain of E10.5 embryos reveal that Z/EG signals in the trigeminal ganglia (*filled arrowheads*) is significantly reduced in *Tmem2^CKO^;Z/EG* embryos (Fig 4B), suggesting that NCCs arriving at these structures is reduced. Sox10 immunostaining confirms that the number of post-migratory NCCs in these structures is indeed reduced in *Tmem2^CKO^;Z/EG* embryos (Fig 4B). Interestingly, *Tmem2^CKO^;Z/EG* embryos exhibit an abnormally folded midbrain neuroepithelium that is Z/EG-positive (*bracket* in Fig 4B), which also indicates aberrant migration of *Tmem2*-deficient NCCs from the dorsal midbrain. In the trunk neural tube, the contribution of Z/EG-positive cells to dorsal root ganglia is reduced in *Tmem2^CKO^;Z/EG* embryos (*arrowheads* in Fig 4C), whereas the domain occupied by Z/EG-positive cells within the dorsal neural tube is expanded in *Tmem2^CKO^;Z/EG* embryos (*bracket* in Fig 4C). Impaired contribution of TMEM2-deficient NCCs to dorsal root ganglia was also demonstrated by immunostaining for Sox10 and counting of Sox10-positive cells (Fig 4C, *Bar graph*). Overall, these results support the idea that *Tmem2* deficiency causes perturbed NCCs emigration from the neural tube and thus results in their reduced contribution to the NCC-derived structures.

**Fig 4.**
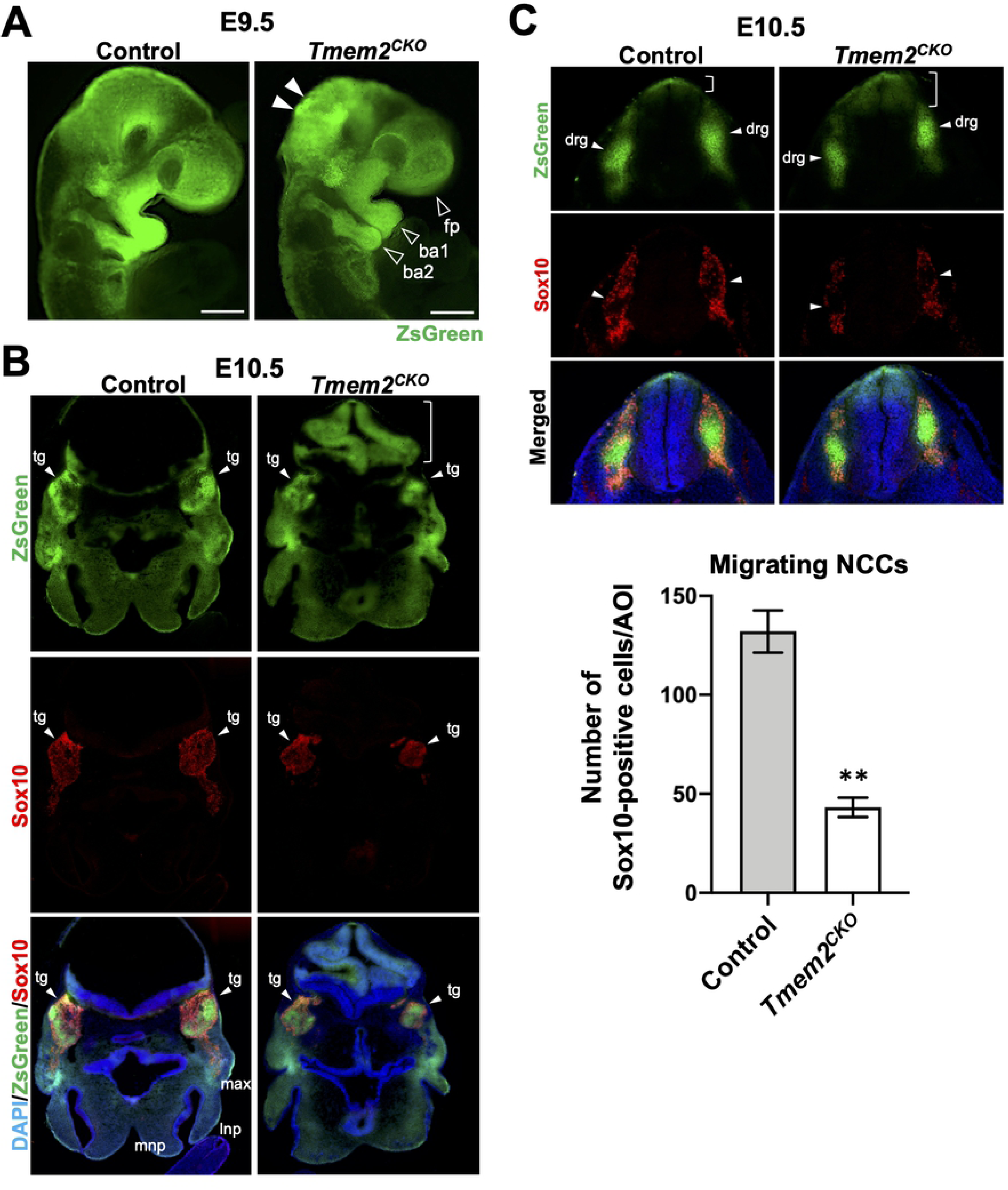
Lineage tracing of NCCs. (**A**) Whole-mount fluorescence images of *Tmem2^CKO^;Z/EG* and control embryos at E9.5. In *Wnt1*-*Tmem2^CKO^;Z/EG* embryos, domains occupied by Z/EG-labeled NCCs are smaller and the intensity of Z/EG signals is lower in the facial prominence, first and second branchial arches (*open arrowheads*). Aberrant accumulation of Z/EG-positive cells was observed in the midbrain and hindbrain of *Tmem2^CKO^;Z/EG* embryos (*filled arrowheads*). (**B**) Transverse sections of *Tmem2^CKO^;Z/EG* and control embryos at E10.5 stained for Sox10. *Arrowheads* indicated trigeminal ganglia (*tg*). The number of Sox10-positive NCCs arrived at the trigeminal ganglia is significantly reduced in *Tmem2^CKO^;Z/EG* embryos. *Tmem2^CKO^;Z/EG* embryos also exhibit abnormal folding of the hindbrain neuroepithelium that contains Z/EG-labeled cells (*bracket*). (**C**) Transverse sections through the caudal neural tube of *Tmem2^CKO^;Z/EG* and control embryos at E10.5 stained for Sox10. In *Tmem2^CKO^;Z/EG* embryos, NCCs migrated to the dorsal root ganglia (*drg*, *arrowheads*) is decreased, while those remaining in the neural tube is increased (*brackets*). Graph shows the quantification of migrating NCCs in *Tmem2^CKO^;Z/EG* and control embryos at E10.5. The number of Sox10-positive cells were counted in transverse sections through the caudal trunk of *Tmem2^CKO^;Z/EG* and control embryos. Means ± SD (*n* = 5) are shown as horizontal bars. *P* values were determined by unpaired Student’s *t*-test. \*\**P* < 0.01. Scale bars, 500 μm in **A**-**C**. *fp*, facial prominence; *ba1*, the first branchial arch; *ba2*, the second branchial arch; *lnp*, lateral nasal process; *max*, maxillary process; *tg*, trigeminal ganglion; *mnp*, medial nasal process; *drg*, dorsal root ganglia.

### *Tmem2*-deficient O9-1 mouse neural crest cells are defective in migration on HA-containing substrates

A considerable amount of evidence indicates that the motility of NCCs is mediated by integrins and the formation of focal adhesions (FAs) [38–42]. While a thin HA coating on glass or plastic can mediate cell attachment, thick HA deposits on the pericellular and extracellular matrices act as a repulsive barrier to cell adhesion and migration [43–46]. This is due to the fact that the gel-like HA matrix, acting as a steric barrier, interferes with the engagement of cell surface integrins with their ECM ligands [35]. The migratory route of NCCs, including the dorsolateral aspect of the neural tube into which early NCCs emigrate, contains high levels of HA [12] (see also Fig 2C). Thus, it is conceivable that NCCs use cell surface TMEM2 to overcome this HA barrier to achieve integrin-mediated FA formation. In this hypothesis, NCCs phenotype of *Tmem2^CKO^* mice steams from the inability of *Tmem2*-deficient NCCs to establish robust integrin interactions with with ECM ligands in the presence of abundant HA.

To address this hypothesis, we used shRNA transfection to generate *Tmem2*-depleted O9-1 cells, a cranial NCCs line [47] (Fig 5A), and examined the effect of *Tmem2* knockdown on O9-1 cell behavior on HA-containing substrates. As shown in Fig 5B, control O9-1 cells degrade substrate-bound HA in a pattern that resembles the distribution of FAs. Immunostaining for vinculin, an FA marker, confirms that control O9-1 cells form FAs at the sites of HA degradation (Fig 5C). *Tmem2*-depleted O9-1 cells, in contrast, exhibit markedly reduced HA degradation. Moreover, vinculin immunostaining reveals that FA formation is greatly attenuated in these cells (Fig 5C). Thus, O9-1 cells employ TMEM2 to mediate HA degradation in the ECM as a means of promoting FA formation.

**Figure 5.**
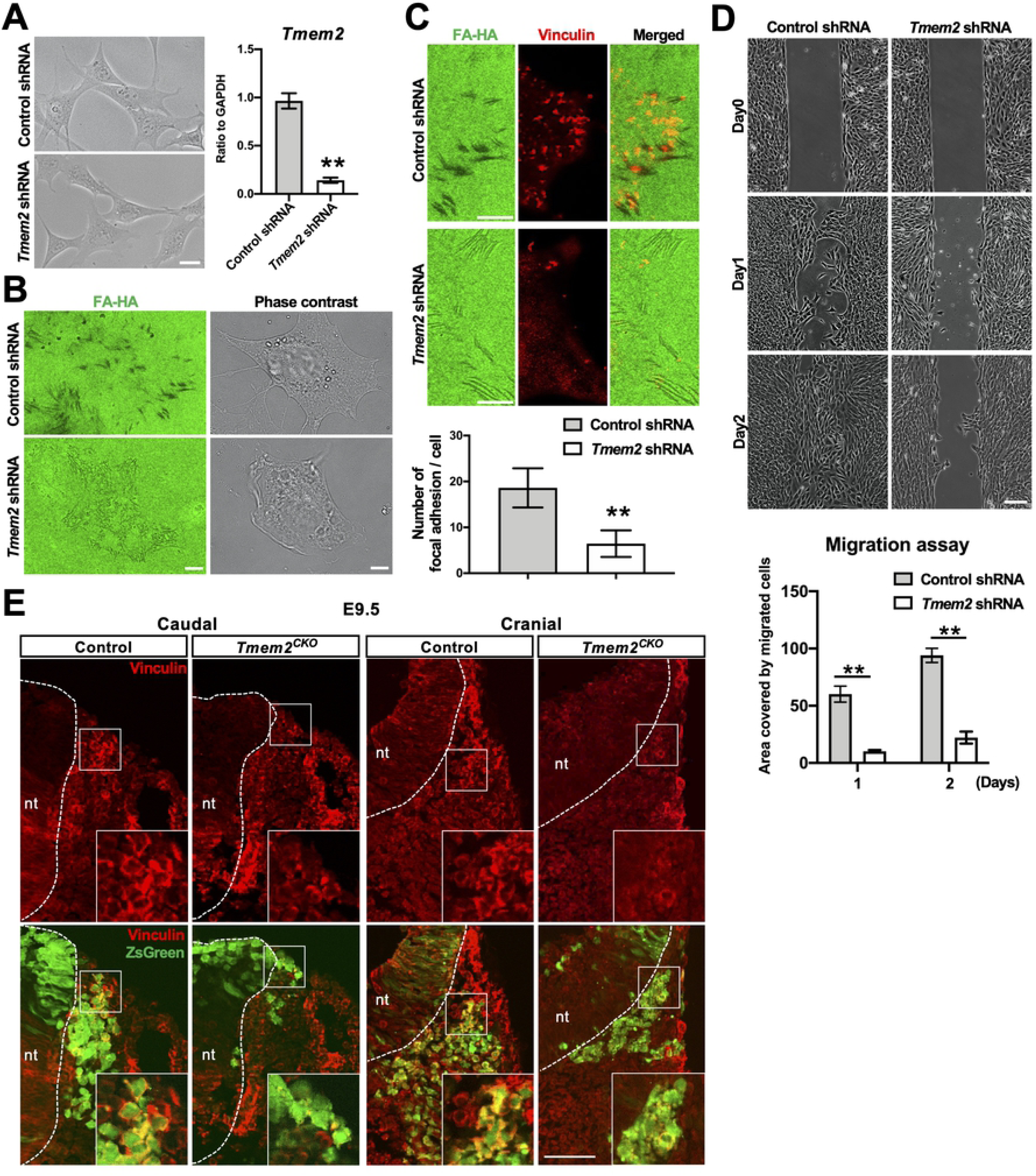
*Tmem2*-deficient NCCs are defective in focal adhesion formation and migration under HA-rich environment. (**A**) Representative images of *Tmem2*-depleted and control O9-1 cells cultured on a regular culture dish. Expression of *Tmem2* is evaluated by qPCR. *Gapdh* was used as an internal control for normalization. Means ± SD (*n* = 5) are shown as horizontal bars. *P* values were determined by unpaired Student’s *t*-test. \*\**P* < 0.01. (**B**) Contact-dependent *in situ* degradation of substrate-immobilized HA by O9-1 cells. Cells were cultured for 16 h on glass coverslips immobilized with fluorescent HA (FA-HA). *In situ* HA degradation activity is detected as dark spots/streaks in FA-HA substrates. Note that wild-type O9-1 cells eliminate substrate-bound HA in a pattern reminiscent of focal adhesions. In *Tmem2*-depleted O9-1 cells, this activity is greatly diminished. (**C**) Colocalization of sites of HA degradation and FAs. *Tmem2*-depleted and control O9-1 cells cultured on Col1/HA mixed substrates were immunostained for vinculin. The number of FAs per cell was quantitatively compared between *Tmem2*-depleted and control O9-1 cells (*bar graph*). Data represent mean ± SD (>30 cells per condition pooled from three independent experiments). *P* values were determined by unpaired Student’s *t*-test. \*\**P* < 0.01. (**D**) Representative images of the migration of *Tmem2*-depleted and control O9-1 cells into a cell-free gap on Col1/HA mixed substrates. Top panels show images of gaps immediately after removal of the ibidi 2-well Culture-Insert. Other panels show images of gaps after a 24 or 48 h incubation. Graph shows the quantitative analysis of cell migration. Data represent the mean ± SD of the gap area covered by migratory cells relative to the area of the original gap (n = 5 per condition). Data are representative of at least three independent experiments. *P* values were determined by two-way ANOVA with Bonferroni’s multiple comparison test. \*\**P* < 0.01. (**E**) Vinculin accumulation in the cortex of migrating NCCs is reduced in *Tmem2*-ablated NCCs *in vivo*. Transverse sections of the neural tube (*nt*) in the caudal and cranial regions of E9.5 *Tmem2^CKO^;Z/EG* and control embryos were stained for vinculin. Boxed areas are enlarged images of the migrating NCCs. nt, neural tube. Scale bars: 5.0 μm in **A**; 2.5 μm in **B**, **C**; 200 μm in **D;** 150 μm in **E**.

To model NCCs emigration into HA-rich tissues surrounding the neural tube, we performed a wound healing-type migration assay on mixed substrates consisting of type I collagen (Col1) and HA (average molecular weight 1200∼1400 kDa). This *in vitro* model allows us to mimic the emigration of leading edge NCCs into the HA-rich interstitial ECM. Control O9-1 cells are highly migratory on these mixed substrates; cells at the boundary of the wound readily migrate into the HA-rich gap (Fig. 5D). In contrast, migration into the gap is significantly reduced in *Tmem2*-depleted O9-1 cells compared with control cells (Fig. 5D). These results demonstrate the functional importance of TMEM2 in migration of O9-1 cells in an HA-rich environment.

To examine the *in vivo* relevance of these findings, we analyzed the localization of vinculin in NCCs of *Tmem2^CKO^;Z/EG* and control (i.e., *Wnt1-Cre;Tmem2^wt/wt^;Z/EG*) embryos at E9.5. In control embryos, individual NCCs emigrating from the neural tube exhibit strong peripheral vinculin immunoreactivity at the periphery of individual cells (Fig 5E), an observation consistent with a previous report [48]. This pattern of subcellular vinculin localization is observed in NCCs in both cranial and caudal regions (Fig 5E). In *Wnt1-Tmem2^CKO^;Z/EG* embryos, in contrast, vinculin immunoreactivity is greatly diminished in emigrating NCCs and little vinculin localization at the cell periphery is observed in these NCCs (Fig 5E). This is consistent with a recent report that formation of the integrin–vinculin–talin complex is necessary for NCCs emigration [49]. Thus, it is likely that reduced NCCs emigration in *Tmem2^CKO^* mice is caused by the impaired NCC’s ability to overcome the barrier effect of HA and form focal adhesion complexes. This further supports the notion that TMEM2-mediated ECM remodeling plays a critical role in the emigration of NCCs and in morphogenesis of NCC-derived tissues.

### *Tmem2* deficiency induces apoptosis in the neural crest-derived tissues

In addition to NCCs emigration defects, *Tmem2^CKO^* embryos exhibit marked impairment in the growth of NCC-derived tissues morphogenesis of NCC-derived tissues (see Fig 1D). To define the mechanisms underlying this phenotype, we analyzed cell proliferation and survival in these tissues. Intriguingly, TUNEL assays reveal that the number of TUNEL-positive cells is markedly increased in the lateral and medial nasal processes and branchial arches of *Tmem2^CKO^* embryos compared to these same areas in control embryos (Fig 6A**, S6A Fig**). Increased in cell death is rather specific for the facial processes and branchial arches; the number of TUNEL-positive cells is not increased in the developing heart of *Tmem2^CKO^* embryos (**S6A Fig**). In contrast to cell death, cell proliferation does not appear to be affected by *Tmem2* ablation; Immunostaining of phospho-histone H3 (PHH3) in these NCC-derived tissues demonstrates no detectable difference in the number of proliferative cells between *Tmem2^CKO^* and control embryos (Fig 6B**, S6B Fig**).

**Figure 6.**
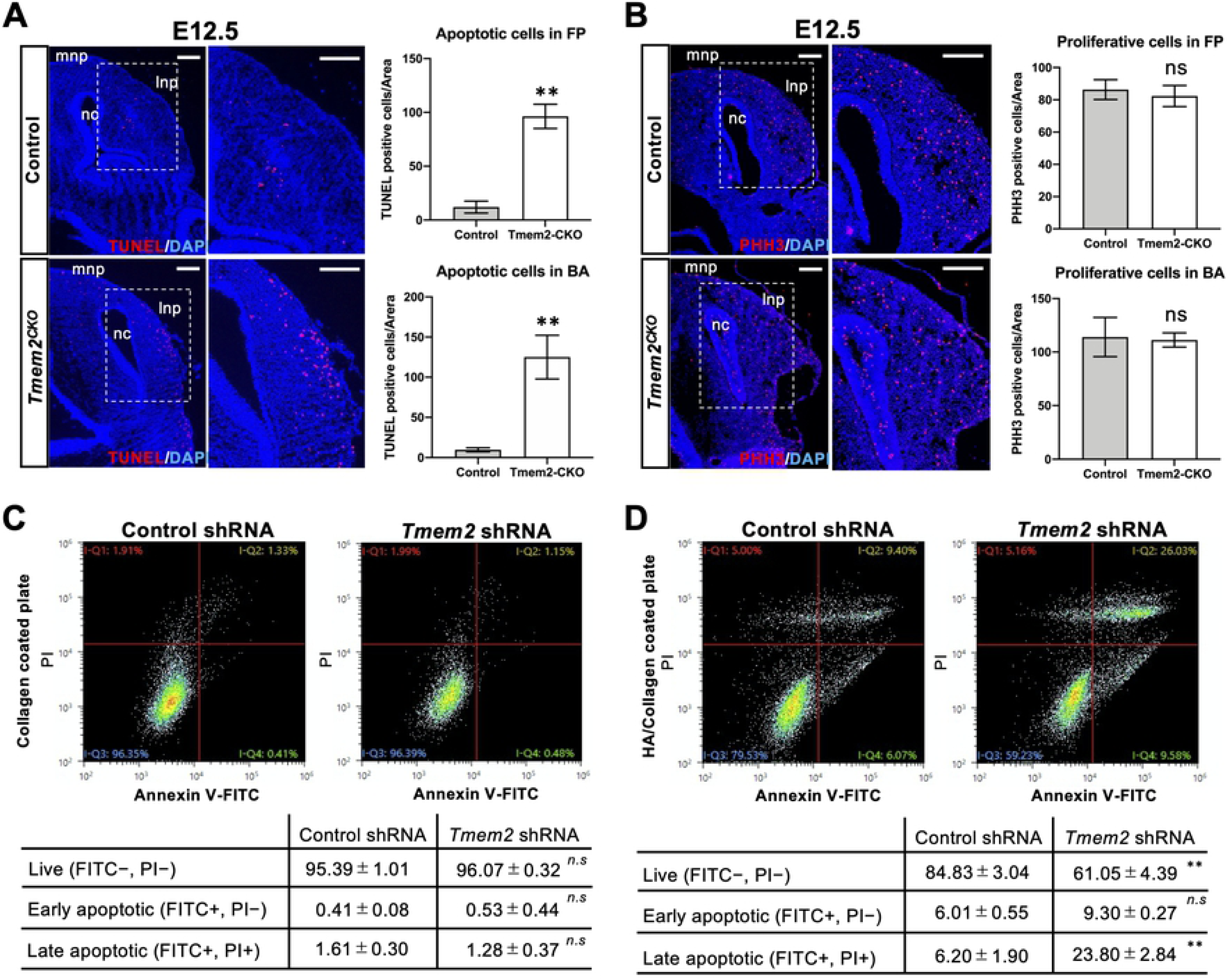
Increased apoptosis of post-migratory NCCs in *Tmem2*^CKO^ embryos. (**A**) Analysis of cell death in the facial processes of E12.5 embryos. Transverse sections through the facial processes were analyzed by TUNEL assays. Graphs on the right represent the number of TUNEL-positive cells per unit area of the facial processes (*top*) and the first branchial arch (*bottom*). Means ± SD (*n* = 3) are shown as horizontal bars. *P* values were determined by unpaired Student’s *t*-test. \*\**P* < 0.01. (**B**) Analysis of cell proliferation in the facial processes of E12.5 embryos. Transverse sections through the facial processes were stained with anti-PHH3 antibody. Graphs on the right represent the number of PHH3-positive cells per unit area of the facial processes (*top*) and the first branchial arch (*bottom*). *mnp*, medial process; *lnp*, lateral nasal process; *nc*, nasal cavity. Means ± SD (*n* = 3) are shown as horizontal bars. *n.s.*, not significant by unpaired Student’s *t*-test. (**C, D**) Analysis of apoptosis of *Tmem2*-depleted and control O9-1 cells cultured on HA-free and HA-containing substrates. *Tmem2*-depleted and control O9-1 cells were cultured for 48 h on Col1 only (**C**) and Col1/HA mixed (**D**) substrates. Cells were then stained with propidium iodide (PI) and anti-annexin V antibody, followed by the analysis of apoptosis by flow cytometry. Lower tables show the quantification of live (lower left quadrant in the charts), early apoptotic (lower right quadrant), and late apoptotic (upper right quadrant) cells. Note that on HA-containing substrates (**D**), significantly greater proportion of Tmem2-depleted O9-1 cells undergo apoptosis than control O9-1 cells. On the other hand, there is no difference between *Tmem2*-depleted and control O9-1 cells on HA-free substrates (**C**). *P* values were determined by unpaired Student’s *t*-test. \*\**P* < 0.01; *n.s.*, not significant. Scale bars: 250 μm.

To examine whether the increased cell death in *Tmem2^CKO^* mice is due to cell autonomous effects of *Tmem2* ablation, we quantitatively analyzed the incidence of apoptosis in O9-1 cells cultured on HA-containing substrates. *Tmem2*-depleted and control O9-1 cells were cultured on Col1 only or on Col1/HA mixed substrates for 48 h. Both depleted and control cells adhere equally well under these conditions. Apoptotic cell death was analyzed by double-staining with anti-annexin V antibody and propidium iodide (PI), followed by flow cytometric analysis of live, early apoptotic, and later apoptotic cells [50]. On Col1 only substrates, there is no difference in the number and pattern of apoptotic (annexin V/PI double-positive) cells between *Tmem2*-depleted and control O9-1 cells (Fig 6C). On the other hand, the number of late apoptotic cells is significantly increased in *Tmem2*-depleted population on Col1/HA mixed substrates (Fig 6D). These *in vitro* data indicate that the apoptotic phenotype of NCC-derived craniofacial components in *Tmem2^CKO^* embryos is a direct consequence of TMEM2 function.

## Discussion

The functional importance of HA in the development of NCC-derived tissues has been demonstrated using genetic models in which hyaluronan synthase genes are mutated or ablated [3, 6, 7]. On the other hand, despite the fact that tissue levels of HA are dynamically regulated by extremely rapid HA turnover, much less is known about the role of HA degradation in embryonic development. The paucity of data related to this issue is largely due to uncertainty regarding the identity of the hyaluronidase(s) responsible for extracellular HA degradation in the developing embryos. In the present study, we demonstrate a critical role for the cell surface hyaluronidase TMEM2 and TMEM2-mediated HA degradation in neural crest development, based on our finding that TMEM2 deficiency leads to defective morphogenesis of NCC-derived structures. Using *Tmem2* knock-in reporter mice reveals that in wild-type embryos, *Tmem2* is expressed in locations involved in NCCs generation and differentiation. These areas include the midline of the neural tube, the branchial arches, the endocardium, and the craniofacial primordia. Consistent with the notion that TMEM2 functions as a hyaluronidase, these regions are generally devoid of HA in wild-type embryos, whereas the same areas in *Tmem2^CKO^* mice exhibit high levels of HA accumulation. NCC-targeted ablation of *Tmem2* results in defective morphogenesis of NCCs derivatives, most notably the craniofacial structure, confirming the essential role of this cell surface hyaluronidase in NCCs development.

Several lines of evidence support the conclusion that these *Tmem2*-dependent defects in NCCs derivatives are due to impaired emigration and migration of *Tmem2*-deficient NCCs. Lineage tracing analysis of *Tmem2^CKO^* embryos reveals accumulation of Sox9-positive pre-migratory cells in the neural tube, while the number of Sox10-positive migratory cranial NCCs is significantly reduced. In addition, our *in vitro* assays using O9-1 mouse neural crest cells demonstrate that *Tmem2* expression is required for NCCs to assemble robust focal adhesions and to migrate on HA-containing substrates (Fig 5). Taken together, these results suggest that temporal and spatial regulation of extracellular HA degradation by TMEM2 is essential for NCCs migration from the neural tube during NCCs development.

We have recently reported that tumor cells use TMEM2 to locally remove matrix-associated HA in the vicinity of adhesion sites, thereby facilitating the formation of mature FAs and associated tumor cell migration, and that this action of TMEM2 is coordinated with integrin function via their direct interactions between the two proteins [30]. This observation has a direct relevance to neural crest development, because the functional importance of integrins and FAs in NCCs emigration and migration has been demonstrated by a number of studies [38-42, 51, 52]. NCCs undergo a process of collective migration, in which they migrate as chains of cells. Cells at the leading edge of collectively migrating NCC chain are polarized, exhibiting distinct leading edges in which integrins are incorporated into FAs that provide links between the actin cytoskeleton and the ECM [41, 53]. In fact, it has recently been shown that inhibition of any of the key FA components, namely integrin-β1, vinculin, or talin, leads to strong impairment of the onset of NCCs migration *in vivo* without significantly affecting cell attachment [38]. We propose that the critical contribution of TMEM2 to NCCs migration is its ability to remove HA that would otherwise sterically impair cell surface integrin interaction with the ECM. In other words, without TMEM2, leading NCCs in the collectively migrating population are not capable of achieving the integrin-ECM engagement needed for robust FA assembly. Consistent with this model, our *in vitro* wound healing assays using *Tmem2*-depleted O9-1 cells shows reduced migration of NCCs from wound borders into at the border of the wounds into HA-containing substrates. Moreover, *Tmem2*-depleted O9-1 cells are not capable of forming FAs on these substrates (Fig 5). We also demonstrate that *Tmem2*-ablated NCCs *in vivo* are incapable of recruiting vinculin in their subcortical areas (Fig 5), supporting the *in vivo* relevance of our model. Taken together, our results strongly suggest that TMEM2-dependent degradation of HA is essential for NCCs emigration due to enhancement of integrin-ECM engagement.

Another notable finding of our current study is the increased apoptosis of post-migratory NCCs in craniofacial tissues of *Tmem2^CKO^* mice (Fig 6A). This *in vivo* finding is further supported by results from *in vitro* experiments with *Tmem2*-depleted O9-1 cells (Fig 6D). There are several potential mechanisms, not necessarily mutually exclusive in nature, by which TMEM2 could be functionally involved in post-migratory NCCs survival. Given the role of TMEM2 to facilitate FA formation and cell adhesion in HA-rich environments [30], it is possible that *Tmem2*-deficiency renders NCCs susceptible to anoikis, a form of programmed cell death that occurs in anchorage-dependent cells [54]. Without TMEM2, NCCs in HA-rich target tissues may not be able to establish the effective cell adhesions needed to avoid anoikis. Another potential mechanism for the increased apoptosis of *Tmem2*-deficient post-migratory NCCs may be elevated endoplasmic reticulum (ER) stress in the absence of TMEM2. In searching for genes needed for cells to survive ER-based stress due to protein misfolding stress, Schinzel et al. [55] identified TMEM2 as a potent protective factor against ER stress damage. Although the precise mechanism by which TMEM2 regulates NCCs survival remains to be determined, these observations collectively suggest that proper expression of TMEM2 is important for the survival of post-migratory NCCs at their destination sites.

In addition to abnormalities in craniofacial tissues, *Tmem2^CKO^* embryos exhibit cardiovascular abnormalities. This observation is consistent with previous reports on zebrafish *tmem2* mutants [26, 27], and also resonates with the fact that mice with mutations in genes involved in HA metabolism and function, including *Has2* (encoding a HA synthase) and *Vcan* (encoding the HA-binding chondroitin sulfate proteoglycan versican), develop cardiovascular abnormalities [3, 56, 57]. Altered levels of HA deposition, due either to abnormal HA accumulation or HA insufficiency, might result in cardiovascular abnormalities based at least partly on defective cranial NCCs development. Further experiments using additional *Cre* deleter lines specific for vascular lineages will help address the roles of TMEM2 in cardiovascular development.

In conclusion, our results demonstrate that the membrane-anchored hyaluronidase TMEM2 plays a critical role in neural crest development. As such, this paper reveals for the first time that the catabolic processing of HA has a specific regulatory role in embryonic morphogenesis, and that its dysregulation leads to severe developmental defects. In this regard, it is noteworthy that a nonsynonymous 1358G>A (C453Y) mutation in exon 5 of the *TMEM2* gene, which is predicted to be pathogenic, has been identified in an individual with hypertelorism, myopia, retinal dystrophy, abnormality of the sternum, joint laxity, and inguinal hernia (ClinVar, NCBI, NIH: accession number: VCV000827846 and VCV000827847). It remains to be seen whether mutations/polymorphisms in *TMEM2* are associated with craniofacial malformation in humans.

## Materials and Methods

### Mice

A conditional *Tmem2* null allele (*Tmem2^flox^*) was created by Cyagen Biosciences (Santa Clara, CA) using TurboKnockout gene targeting methods. Mouse genomic fragments containing homology arms and the conditional knockout region were amplified from a BAC library and were sequentially assembled into a targeting vector together with recombination sites and selection markers. The targeting vector was electroporated into C57BL/6-derived ES cells, followed by drug selection and isolation of drug-resistant clones. The resultant *Tmem2^flox^* allele contains two loxP sites flanking exons 4 and 5 of the *Tmem2* gene, so that these two exons will be deleted by Cre-mediated recombination. Exon 5 harbors amino acid residues (R265, D273, D286), mutagenesis of any of which abrogates the hyaluronidase activity of TMEM2 [28]. Moreover, conjugation of exon 3 to exon 6 due to the deletion of exons 4 and 5 results in a frameshift and a premature stop codon. Mice carrying homozygous *Tmem2^flox^* alleles are developmentally normal and fertile, confirming that the non-recombined *Tmem2^flox^* allele is fully functional. Mice with conditional *Tmem2* ablation targeted to NCCs were generated by crossing the *Tmem2^flox^* allele and the *Wnt1-Cre* transgene [34]. *Wnt1-Cre* mice in a C57BL/6 background were obtained from Prof. Sachiko Iseki of Tokyo Medical and Dental University. Resultant *Wnt1-Cre;Tmem2^flox/wt^* male mice were mated with *Tmem2^flox/flox^* or *Tmem2^flox/wt^* female mice to obtain *Tmem2* conditional knockout mice (i.e., *Wnt1-Cre;Tmem2^flox/flox^*). Littermates inheriting an incomplete combination of the above alleles were used as controls (referred to as wild type). Genotyping of mice and embryos was performed by PCR with the specific primers listed in supplemental table 1 (**S1 Table**), using DNA prepared from tail biopsies and yolk sacs. For *in vivo* fate mapping experiments, ZsGreen reporter mice (R26;ZsGreen; JAX mice 007906) were purchased from The Jackson Laboratory (Bar Harbor, ME). Mice with *Tmem2^flox^* allele and the R26;ZsGreen transgene (Z/EG) were crossed to create *Tmem2^flox/wt^;Z/EG* mice*. Wnt1-Cre;Tmem2^flox/wt^* male mice were mated with *Tmem2^flox/wt^;Z/EG* female mice to obtain ZsGreen reporter mice with *Tmem2* conditional knockout (*Tmem2^flox/flox^;Wnt1-Cre;Z/EG*) or control allele (*Tmem2^wt/wt^;Wnt1-Cre;Z/EG*). Primer sequences for genotyping are listed in **S1 Table**.

### Creation of the Tmem2-FLAG knock-in allele

Pronuclear stage embryos from C57BL6/J mice were purchased from ARK Resource (Kumamoto, Japan). Recombinant Cas9 protein, crRNA and tracrRNA were obtained from Integrated DNA technology. Single-stranded oligodeoxynucleotides for insertion of a FLAG epitope were designed with 30 bp sequence homology on each side of Cas9-mediated double strand break (see **S2 Table for sequence information**). For generation of *Tmem2*-FLAG knock-in (*Tmem2-FLAG^KI^*) mice, we used the Technique for Animal Knockout System by Electroporation (TAKE) [58]. Briefly, embryos were washed twice with Opti-MEM solution and aligned in the electrode gap filled with 50 μl of Cas9/gRNA(crRNA-tracrRNA complex)/ssODN (200/100/100 ng/μl) mixture. The intact embryos were subjected to electroporation using poring (225V) and transfer pulses (20V). After electroporation, embryos were returned to KSOM media at 37°C. Genome edited 2-cell embryos were transferred to oviducts of pseudopregnant ICR female mice, and genomic DNA from newborn mice was analyzed by PCR (see **S1 Table** for primer sequences).

### Immunohistochemistry, TUNEL staining, and *in situ* hybridization

Embryos were fixed with 4% PFA and incubated at 4°C overnight with DAPI (4′,6-diamidino-2-phenylindole) diluted 1:1000 (Dojindo, **#**D523-10). Immunostaining of frozen sections was performed as previously described [59]. The following antibodies were used in this study: mouse monoclonal anti-FLAG (F9291, 1:200), rabbit polyclonal anti-FLAG (F7425, 1:200), mouse monoclonal anti-vinculin (V4505, 1:200), and mouse monoclonal α-SMA (A5228, 1:200) from Sigma; rabbit polyclonal anti–SOX9 (#82630, 1:100) from Cell Signaling Technology; rabbit polyclonal anti-Sox10 (ab27655, 1:100) from abcam; mouse monoclonal anti-phospho-histone H3 (PHH3) (05-746R, 1:200) from MilliporeSigma; Alexa 488-labelled goat anti-mouse IgG (A28175, 1:500), Alexa 488-labelled goat anti–rabbit IgG (#A11034, 1:500), Alexa 555-labelled goat anti-mouse IgG (A21422, 1:500) and Alexa 555-labeled goat anti-rabbit IgG (A21428, 1:500), Alexa 555-labeled goat anti-rat IgG (A21434, 1:500), Alexa 555-labeled streptavidin (S32355, 1:500) from Invitrogen; horseradish peroxidase-conjugated goat anti-rabbit IgG (1706565, 1:500) and goat anti-mouse IgG (1706516, 1:500) from Bio-Rad. DAPI were purchased from Invitrogen. Apoptotic cells were identified by using an *in situ* cell death detection kit (Roche, catalog #11684795910) according to the manufactureŕs instructions. Whole mount nuclear fluorescence imaging was utilized to capture embryonic morphological features. Biotinylated HA binding protein (bHABP) was used for detection of HA as previously described [60]. Whole-mount *in situ* hybridization was performed as described previously [61]. The digoxigenin-labelled antisense RNA probes were produced using a DIG RNA labeling kit (Roche, #11277073910) according to the manufacturer’s protocol. A minimum of three embryos of each specimen type were examined per probe.

### Immunoprecipitation and immunoblotting

Protocols for immunoprecipitation and immunoblotting were described previously [62]. Briefly, cells were lysed in ice-cold 50 mM Tris-HCl (pH 7.5) containing 250 mM NaCl, 0.1% Triton X-100, 1 mM EDTA, 50 mM NaF, 0.1 mM Na_3_VO_4_, 1 mM DTT, 0.1 mM leupeptin, 0.1 μg/mL soybean trypsin inhibitor, 10 μg/mL L-1 chloro-3-(4-tosylamido)-4 phenyl-2-butanone (TPCK), 10 μg/mL L-1 chloro-3-(4-tosylamido)-7-amino-2-heptanone-hydrochloride (TLCK), 10 μg/mL aprotinin, and 50 μg/mL phenylmethylsulfonyl fluoride (PMSF). Following a 30 min lysis period on ice, samples were centrifuged at 15,000 rpm for 20 min at 4°C to prepare cell lysates. Ten μg of lysate was subjected to SDS-PAGE on an 8–16% Tris-glycine gel (ThermoFisher), followed by electroblotting onto an Immobilon PVDF membrane (EMD Millipore). Rabbit polyclonal antibody against TMEM2 was purchased from Sigma Aldrich (SAB2106587). The ECL Western Blotting Substrate (Nacalai Tesque, #07880) was used to detect signals. Immunoprecipitation was performed with Dynabeads streptavidin magnetic beads and biotinylated anti-FLAG antibody (MilliporeSigma, #F9291) as previously described [63], according to the manufacturer’s instructions. Briefly, for detecting endogenous C-terminal FLAG-tagged TMEM2, cell lysates were obtained by treating embryos with lysis buffer containing 1% Brij 96, 25 mM HEPES (pH 7.5), 150 mM NaCl, and 5 mM MgCl_2_. Wild-type embryos were used as a negative control for the experiment. Dynabeads streptavidin magnetic beads (Invitrogen, DB65801) were washed 3 times with wash buffer and incubated with the biotinylated anti-FLAG antibody in PBS for 30 min at room temperature using gentle rotation. The antibody-coated streptavidin magnetic beads were separated with a magnet and the coated beads were washed 4–5 times in PBS containing 0.1% BSA. Washed antibody-coated streptavidin magnetic beads were added to the cell lysates and incubated overnight at 4°C. Antibody-coated streptavidin magnetic beads were captured with a magnet and washed 2–3 times. Immunoprecipitates solubilized with elution buffer were subjected to Western blotting.

### Quantification of HA

Neural crest-derived craniofacial tissues were dissected from *Tmem2^CKO^* and control embryos at E12.5. Following incubation in ice-cold lysis buffer for 30 min as described above, samples were centrifuged at 15,000 rpm for 20 min at 4°C to prepare tissue lysates. Tissue HA contents were measured by latex-sensitized immunoturbidimetry (Hyaluronic Acid LT Assay, Fujifilm Wako Chemicals GmbH, Neuss, Germany) using a Hitachi 917s.

### Culture and lentiviral transduction of O9-1 cells

The O9-1 mouse cranial neural crest cell line was purchased from MilliporeSigma (SCC049) and cultured in complete ES cell medium containing 15% FBS and LIF (MilliporeSigma, ES-101-B) with 100 units/ml penicillin-streptomycin (Invitrogen) at 37 °C in a humidified atmosphere containing 5% CO_2_ according to the manufacturer’s instructions. To knockdown *Tmem2* expression in O9-1 cells, we used lentivirus-mediated shRNA transduction. Lentivirus particles targeting mouse *Tmem2* (Mission shRNA, TRCN0000295501) and their controls (Mission shRNA, SHC005) were purchased from MilliporeSigma. Lentivirus particles were added to O9-1 cells cultured in growth media supplemented with 5 μg/ml polybrene and cultured for 2 days. Cells transduced with lentiviral shRNAs were selected and maintained in the presence of 10 μg/mL puromycin.

### RNA extraction and qPCR analysis

Neural crest-derived craniofacial tissues were dissected from *Tmem2^CKO^* and control embryos at E12.5. Total mouse RNA was isolated using RNeasy mini kit (Qiagen) and reverse transcribed to cDNA using an oligo (dT) with reverse transcriptase (Takara Bio) as previously described [63]. For real-time PCR, aliquots of total cDNA were amplified with the TaqMan Fast Universal PCR Master Mix (Applied Biosystems, Foster City, CA). Data acquisition and analysis were performed with a Step One Real-Time PCR System using Step One Software, Version 2.1 (Applied Biosystems). The PCR products were quantified using *gapdh* as the reference gene. The primers and TaqMan probes were purchased from Applied Biosystems.

### *In situ* HA degradation assay

This assay was performed as described previously [30]. Briefly, coverslips were coated with a mix of type I collagen and fluorescein-labeled HA (FAHA-H1, Iwai Chemicals). Trypsinized cells were seeded onto the coated coverslips in 24-well plates at a density of 5 × 10^4^ cells per coverslip and then incubated for 16 h at 37 °C in a CO_2_ incubator. Cells were fixed with 4% paraformaldehyde in PBS for overnight at 4°. In some experiments, fixed cells were permeabilized with PBS containing 0.2% Triton X-100 for 10 min and blocked with PBS containing 1% IgG-free bovine serum albumin (BSA) (Sigma, A2058) for 30 min at RT, followed by immunocytochemistry with mouse monoclonal anti-vinculin antibody (1:200 dilution in 1% BSA–PBS) (Sigma; V9264, clone hVIN-1) and Alexa 555–labelled goat anti–mouse IgG (#A21422, 1:500 dilution in 1% BSA–PBS). DAPI (1:500, Dojindo) was used for nuclear staining and the sections were mounted with fluorescence mounting medium (Dako). Fluorescent images were captured using an all-in-one fluorescence microscope (BZ-X700, Keyence, Osaka, Japan).

### Cell migration assay

A wound healing-type assay was performed using 2-well culture inserts (ibidi; 80209) to create a defined 500 μm cell-free gap on Col1/HA substrates. Glass coverslips were coated with type I collagen and HA as described above. After drying, 2-well culture inserts (ibidi; 80209) were attached to coated coverslips. Insert-attached coverslips were then transferred into 24-well plates, and on the outside, the inserts were filled with PBS. Cells (1 × 10^4^ cells in 70 μl of culture medium per insert) were seeded into the wells and cultured for 2 days. Two days later, culture inserts were detached from coverslips, and coverslips were transferred into new 24-well plates with fresh culture medium. At 24 h and 48 h in culture, phase contrast images were captured by an all-in-one fluorescence microscope (BZ-X700, Keyence, Osaka, Japan). The area in a gap (0.5 mm in distance) between two wells covered by migrating cells was analyzed by ImageJ 1.51s (NIH).

### Image analysis of NCCs migration

For the quantification of pre-migratory and migrating NCCs, transverse sections of the neural tube at the corresponding truncal level were prepared from *Tmem2^CK^*^O^ and control embryos at E9.0. For the analysis of pre-migratory NCCs, sections were stained with anti-Sox9 antibody and DAPI, as described above. The neural tube was outlined in DAPI labeled images, which was defined as a region of interest (ROI). For the analysis of migrating NCCs, sections were stained with anti-Sox10 antibody and DAPI, and a square area (1200 × 1200 μm in Fig. 3C; 2000 × 2000 μm in Fig. 4C) in the dorsal edge of the neural tube was selected as an ROI. In both analyses, the number of labeled cells in each ROI is counted at least 5 consecutive sections per embryo in a total 3-5 embryos by ImageJ 1.51s.

### Statistical analysis

Statistical methods were not used to predetermine sample size. Statistical analyses were performed with GraphPad Prism 8. Student’s two-sided *t* test and two-way ANOVA were used under the assumption of normal distribution and observance of similar variance. A *p* value of <0.05 was considered significant. Bonferroni post hoc analysis was performed where applicable. Values are expressed as mean ± SD. Data shown are representative images; each analysis was performed on at least three mice per genotype. Immunostaining was performed at least in triplicate.

### Ethics Statement

All animal experiments were performed in accordance with the guidelines of the Animal Care and Use Committee of the Osaka University Graduate School of Dentistry, Osaka, Japan. The Committee on the Ethics of Animal Experiments of the same university approved the study protocol. Approval number: 04263, 29-024-0.

## Acknowledgements

We thank Dr. Sachiko Iseki (Tokyo Medical and Dental University) for providing *Wnt1-Cre* mice. We thank Ms. Yuriko Nogami and Ms. Yuki Okamoto for the excellent care and maintenance of our mouse colony and for valuable assistance in the histological, molecular and protein work.

## Supporting information

**S1 Fig. Expression of *Tmem2* in wild-type embryos at E9.5 and E10.5.**

Whole-mount *in situ* hybridization images of *Tmem2* at E9.5 and E10.5. *ba*, branchial arch; *h*, heart; *tg*, trigeminal ganglia; *fp*, facial prominence; *drg*, dorsal root ganglia; *fb*, forebrain; *mb*, midbrain; *hb*, hindbrain. Scale bar: 500 μm.

**S2 Fig. *Tmem2^CKO^* embryos exhibits abnormalities in the neural tube.**

Images show the dorsal views of *Tmem2^CKO^* and control embryos at E12.5. A fraction (4 of 42, 9.5%) of *Tmem2^CKO^* embryos exhibit defects in the neural tube, including incomplete neural tube closure (*filled arrowheads*) and kinking of the neural tube (*open arrowheads*). Scale bars: 500 μm.

**S3 Fig. *Tmem2^CKO^* embryos exhibit abnormal endocardial cushion formation and endocardial cell migration in the outflow tract region.**

Transverse sections through the outflow tract region of E12.5 *Tmem2^CKO^* and control embryos were stained with anti-CD31 (*green*), anti-αSMA (*red*), and DAPI (*blue*). *Asterisk* indicates abnormal aggregates of endocardial cells in endocardial cushion mesenchyme. *Arrowheads* in the *Tmem2^CKO^* embryo point to lack of normal endocardial layer overlying the conotruncal endocardial cushions. Insets show enlarged images of the endocardial cushion mesenchyme. *Ao*, aorta; *ec*, endocardial cushion; *v*, ventricle. Scale bars: 200 μm.

**S4 Fig. Aberrant HA accumulation in the mandible and heart of E12.5 *Tmem2^CKO^* embryos.**

Transverse sections of the mandible and the heart were stained with H&E or double-labeled with bHABP (*red*). Aberrant HA accumulation is observed in the mandible and the heart (in which areas indicated by rectangles are enlarged). *Asterisks* indicate endocardial cushion mesenchyme. Scale bars: 250 μm.

**S5 Fig. Generation of *Tmem2-FLAG^KI^* reporter mice.**

(**A**) Schematic diagram of FLAG-tagged TMEM2 protein expressed from the *Tmem2-FLAG^KI^* locus. (**B**) Expression of TMEM2-FLAG protein in *Tmem2-FLAG^KI^* mice. TMEM2-FLAG protein was immunoprecipitated from the lysate of E11.0 whole embryos with anti-FLAG M2 antibody (*IP*). Precipitated materials were subjected to immunoblotting analysis (*IB*) with anti-TMEM2 antibody.

**S6 Fig. Analysis of apoptosis and proliferation of post-migratory NCCs in the branchial arches and the heart.**

(**A**) Apoptotic cells were detected using TUNEL staining. *Tmem2^CKO^* embryos at E12.5 show the increased number of TUNEL-positive cells in the branchial arches (BA). *Asterisk* indicates the outflow tract region. (**B**) Transverse sections of *Tmem2^CKO^* and control embryos at E12.5 were immunohistochemically stained with anti-PHH3 antibody and DAPI. Apoptosis is increased in the branchial arch of *Tmem2^CKO^* embryo compared to control embryos, whereas cell proliferation is not altered. Scale bars: 250 μm.

**S1 Table. Primer sequences for PCR genotyping of mice.**

**S2 Table. Sequences of crRNA and the inserted FLAG-encoding DNA fragment used for the creation of the *Tmem2-FLAG* knock-in allele (*Tmem2-FLAG^KI^*).**

